# 3CAC: improving the classification of phages and plasmids in metagenomic assemblies using assembly graphs

**DOI:** 10.1101/2021.11.05.467408

**Authors:** Lianrong Pu, Ron Shamir

## Abstract

**Motivation:** Bacteriophages and plasmids usually coexist with their host bacteria in microbial communities and play important roles in microbial evolution. Accurately identifying sequence contigs as phages, plasmids, and bacterial chromosomes in mixed metagenomic assemblies is critical for further unravelling their functions. Many classification tools have been developed for identifying either phages or plasmids in metagenomic assemblies. However, only two classifiers, PPR-Meta and viralVerify, were proposed to simultaneously identify phages and plasmids in mixed metagenomic assemblies. Due to the very high fraction of chromosome contigs in the assemblies, both tools achieve high precision in the classification of chromosomes but perform poorly in classifying phages and plasmids. Short contigs in these assemblies are often wrongly classified or classified as uncertain.

**Results:** Here we present 3CAC, a new three-class classifier that improves the precision of phage and plasmid classification. 3CAC starts with an initial three-class classification generated by existing classifiers and improves the classification of short contigs and contigs with low confidence classification by using proximity in the assembly graph. Evaluation on simulated metagenomes and on real human gut microbiome samples showed that 3CAC outperformed PPR-Meta and viralVerify in both precision and recall, and increased F1-score by 10-60 percentage points.

**Availability:** The 3CAC software is available on https://github.com/Shamir-Lab/3CAC.

**Supplementary information:** Supplementary data are available at *Bioinformatics* online.

## 1 Introduction

The metagenomes of microbial communities are mainly composed of bacterial chromosomes and the associated extrachromosomal mobile genetic elements (eMGEs), such as plasmids and bacteriophages (phages). These eMGEs carry genes related to antibiotic resistance (Calero-Cáceres *et al*., 2019; Wein *et al*., 2019; Lopatkin *et al*., 2017), virulence factors (Kraushaar *et al*., 2017; Sarowska *et al*., 2019) and auxiliary metabolic pathways (Hurwitz and U’Ren, 2016; Rosenwasser *et al*., 2016; Kieft *et al*., 2020). They can frequently move between species in the microbial community (Sitaraman, 2018; Frost *et al*., 2005) and enable their hosts to rapidly adapt to environmental changes (Thomas and Nielsen, 2005; Smalla *et al*., 2015). Despite their important roles in horizontal gene transfer events and in antibiotic resistance, our understanding of these eMGEs is still limited. Part of the difficulty is the challenge of identifying such elements efficiently from mixed metagenomic assemblies (Arredondo-Alonso *et al*., 2017; Krishnamurthy and Wang, 2017; Antipov *et al*., 2019, 2020; Pellow *et al*., 2021; Suzuki *et al*., 2019; Yahara *et al*., 2021).

Multiple algorithms have been developed for identifying either phages or plasmids from metagenomic assemblies in recent years. VirSorter and VirSorter2 identify viral metagenomic fragments by searching for reference homologs and testing enrichment of virus-like proteins (Roux *et al*., 2015; Guo *et al*., 2021). These knowledge-based tools have high precision in virus classification but poor ability to identify novel viruses, due to reference database-associated bias. Other tools, such as DeepVirFinder (Ren *et al*., 2020), Seeker (Auslander *et al*., 2020), and VIBRANT (Kieft *et al*., 2020), use machine learning to learn k-mer signatures of viral sequences and perform better on novel virus classification, since they are more loosely linked to annotation databases. cBar is the first tool designed primarily for plasmid identification in metagenomes (Zhou and Xu, 2010). More recently, two supervised-learning approaches, PlasFlow (Krawczyk *et al*., 2018) and PlasClass (Pellow *et al*., 2020), were shown to classify plasmid fragments better from metagenomic assemblies. Although both phages and plasmids are commonly found in the metagenomes of microbial communities, all of these tools identify either only phages or only plasmids from metagenomic assemblies.

Currently, only two published tools, PPR-Meta (Fang *et al*., 2019) and viralVerify (Antipov *et al*., 2020), can identify phages and plasmids simultaneously from metagenomic assemblies. However, due to the overwhelming abundance of chromosome fragments in the assemblies (usually *≥* 70%), both tools achieve high precision in chromosome classification but very low precision in classification of phages and plasmids (Fang *et al*., 2019; Antipov *et al*., 2020). Moreover, classification of short contigs is challenging for all the existing classifiers, as they analyze each contig independently (Antipov *et al*., 2020; Fang *et al*., 2019; Krawczyk *et al*., 2018; Roux *et al*., 2015; Ren *et al*., 2017). Here we present 3CAC (3-Class Adjacency based Classifier), an algorithm that employs existing two-class and three-class classifiers to generate an initial three-class classification with high precision, and then improves the classification of short contigs and of contigs classified with lower confidence by taking advantage of classification of their neighbors in the assembly graph. Evaluation on simulated and real metagenome datasets with short and long reads showed that 3CAC improved both precision and recall, and increased F1-score by at least 10 percentage points.

## 2 Methods

3CAC accepts as input a set of contigs and its associated assembly graph, uses the classification result of existing tools as a starting point, and repeatedly improves the classification using the assembly graph. Its output is a classification of each contig in the input as phage, plasmid, chromosome, or uncertain. Figure 1 shows the workflow of 3CAC algorithm. The details of the algorithm are described below.

**Fig. 1.**
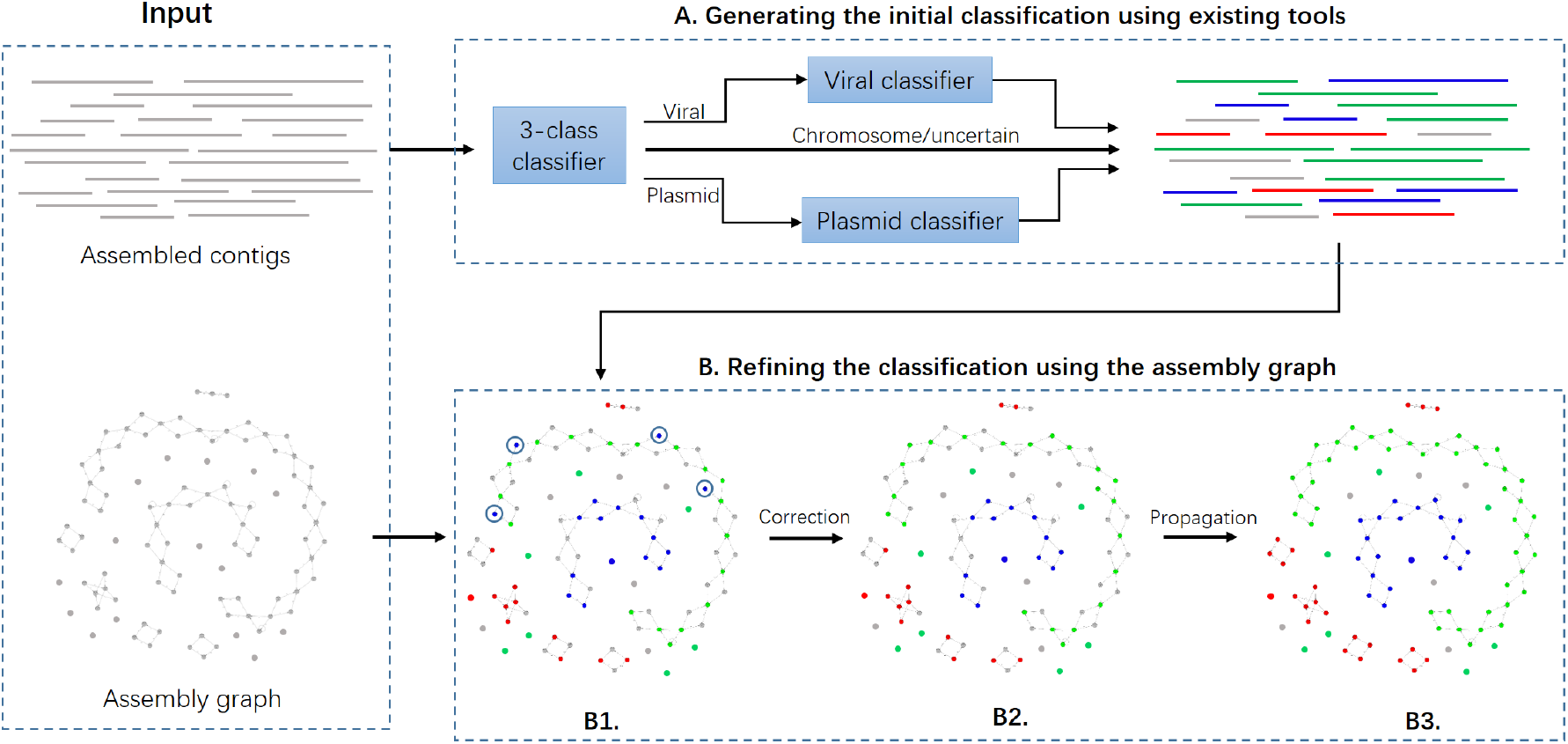
Workflow of 3CAC algorithm. A. Generating the initial classification using existing classifiers. The current version of 3CAC employs PPR-Meta or viralVerify as the 3-class classifier, DeepVirFinder as the viral classifier, and PlasClass as the plasmid classifier. B. Refining the classification using the assembly graph. Contigs/vertices with color red, blue, green, and grey represent contigs classified as phages, plasmids, chromosomes, and uncertain, respectively. (B1) The result of the first phase projected on the assembly graph. (B2) After the correction step. The four contigs encircled in (B1) were corrected. (B3) After the propagation step.

### 2.1 Generating the initial classification

3CAC exploits existing two-class and three-class classifiers to generate an initial three-class classification as follows.

#### (1) Generating a three-class classification

The algorithm runs either viralVerify or PPR-Meta on the set of the input contigs and classifies each contig as phage, plasmid, chromosome, or uncertain. viralVerify was designed to classify contigs as viral, non-viral or uncertain. Moreover, for non-viral contigs, viralVerify can further classify them as plasmid or non-plasmid using *-p* option. Here, we used *-p* option of viralVerify to classify each of the input contigs as viral, plasmid, chromosome, or uncertain. PPR-Meta calculates three scores representing the probabilities of a contig to be classified as a phage, plasmid, or chromosome. By default, PPR-Meta classifies a contig into the class with the highest score. If a specified score threshold is provided and no score passes the threshold, the sequence will be classified as uncertain. Here, we ran PPR-Meta with a score threshold of 0.7.

#### (2) Improving plasmid classification

To improve the precision of plasmid classification, PlasClass is run on contigs classified as plasmids in step (1). PlasClass outputs for each contig the probability that it originated from a plasmid. By default, PlasClass classifies a contig as plasmid if it has a probability > 0.5 and as chromosome otherwise. To assure high precision, here we identify contigs with probability ≥ 0.7 as plasmids. Contigs with probability ≤ 0.3 are moved to the chromosome class. The remaining contigs are reclassified as uncertain.

#### (3) Improving phage classification

Similarly, in order to improve the precision of phage classification, we run DeepVirFinder on all contigs classified as phages in step (1). DeepVirFinder generates a score and a p-value for each input contig. Contigs with higher scores or lower p-values are more likely to be viral sequences. Here, a contig is kept in the phage class if its p-value ≤ 0.03 and moved to the chromosome class if its p-value > 0.03 and its score ≤ 0.5. The remaining contigs are reclassified as uncertain.

We will denote the algorithm up to this step **Initial(vV)** and **Initial(PM)** if viralVerify or PPR-Meta were used in step (1), respectively.

### 2.2 Refining the classification using the assembly graph

In genomics and metagenomics, assembly graphs, such as de Bruijn graphs (Lin *et al*., 2016; Pevzner *et al*., 2001) and string graphs (Myers, 2005; Simpson and Durbin, 2012), are used as the core data structure to combine overlapped reads (or k-mers) into contigs. Initially, nodes are *k*-mers and edges represent (*k* + 1)-long overlaps between them. Each longest path of nodes with indegree and outdegree=1 is then collapsed into a single node representing the corresponding sequence contig. In our description below, nodes in the graph are the contigs, and the neighbors of a contig are its adjacent nodes. Existing classifiers take contigs as input and classify each of them independently based on its sequence. The overlap information between neighboring contigs in the assembly graphs was ignored by all the existing classifiers. However, recent studies showed that neighboring contigs in an assembly graph are more likely to come from the same taxonomic group (Barnum *et al*., 2018; Mallawaarachchi *et al*., 2020). Based on this insight, here we exploit the assembly graph to improve the classification by the following two steps.

#### (1) Correction of classified contigs

A classified contig is called *incongruous* if it has ≥ 2 classified neighbors and all of them belong to same class, while this contig belongs to a different class. We reason that an incongruous contig was wrongly classified and its classification needs to be consistent with its classified neighbors. Therefore, 3CAC scans all the incongruous contigs in decreasing order of the number of their classified neighbors and corrects the classification of each incongruous contig to match its classified neighbors. Note that once an incongruous contig is corrected, this contig and all its neighbors will not be corrected anymore.

#### (2) Propagation from classified contigs to unclassified contigs

An unclassified contig is called *implied* if it has one or more classified neighbors and all of them belong to same class. 3CAC dynamically maintains a sorted list of implied contigs in decreasing order of the number of their classified neighbors. At each iteration, 3CAC classifies the first implied contig on the list according to its classified neighbors and then updates the sorted list. Note that only the unclassified neighbors of the first implied contig need to be updated at each iteration. We repeat this step until the list is empty.

Figure 1B shows the result of applying steps (1) and (2) in a small assembly graph, which is part of the graph generated by assembling simulated long reads (Sim4; see details in the Results section).

We will use the names **3CAC(vV)** and **3CAC(PM)** for the full 3CAC algorithms initialized with viralVerify and PPR-Meta solutions, respectively.

## 3 Results

We tested 3CAC on both simulated and real metagenome assemblies and compared it to PPR-Meta and viralVerify.

### 3.1 Evaluation criteria

3CAC, viralVerify and PPR-Meta were evaluated based on precision, recall, and F1 score, calculated as follows.

- **Precision:** the fraction of correctly classified contigs among all classified contigs. Note that uncertain contigs were not included in the calculation.
- **Recall:** the fraction of correctly classified contigs among all contigs.
- **F1 score:** the harmonic mean of the precision and recall, which can be calculated as: *F* 1 *score* = (2 *\* precision * recall*)*/*(*precision* + *recall*).

Following (Pellow *et al*., 2020; Fang *et al*., 2019), the precision, recall, and F1 score here were calculated by counting the number of contigs and did not take into account their length. The precision and recall were also calculated separately for phage, plasmid and chromosome classification. For example, the precision of phage classification was calculated as the fraction of correctly classified phage contigs among all contigs classified as phages, and the recall of phage classification was calculated as the fraction of correctly classified phage contigs among all phage contigs.

### 3.2 Performance on simulated metagenome assemblies

We generated two short-read and two long-read metagenome assemblies as follows. Sequences of complete bacterial genomes were randomly selected from the NCBI database along with their associated plasmids. The abundance of bacterial genomes was modeled by the log-normal distribution and the copy numbers of plasmids were simulated by the geometric distribution as in (Pellow *et al*., 2020). The phage genomes and their abundance profiles were sampled from (Ren *et al*., 2017). Two metagenomic datasets of different complexities were designed. For each of the datasets, 150bp-long short reads were simulated from the genome sequences using InSilicoSeq (Gourlé *et al*., 2019) and assembled by metaSPAdes (Nurk *et al*., 2016). Long reads were simulated from the genome sequences using NanoSim (Yang *et al*., 2017) and assembled by metaFlye (Kolmogorov *et al*., 2020). The error rate of long reads was 9.8% and their average length was 14.9kb. For each assembly, contigs were matched to the reference genomes used in the simulation by minimap2 (Li, 2018). Contigs having matches to a reference genome with ≥ 90% mapping identity along ≥ 80% of the contig length were assigned to the class of that reference, and these assignments were used as the gold standard to test the classifiers. Table 1 presents a summary of the simulated metagenome assemblies.

**Table 1.** Properties of the simulated and the real metagenome datasets and of their assemblies. The number of genome references for the real human gut metagenomes is the number of all complete chromosome, plasmid and phage genomes in NCBI database.

| Dataset | Read type | Number of reads | # of genome references |  |  | # of assembled contigs |  |  | Short contigs (< 1kb) |
| --- | --- | --- | --- | --- | --- | --- | --- | --- | --- |
|  |  |  | chromosome | plasmid | phage | chromosome | plasmid | phage |  |
| Sim1 | MiSeq | 61M | 50 | 193 | 200 | 12,494 | 1,699 | 696 | 8,991 |
| Sim2 | MiSeq | 100M | 100 | 410 | 500 | 40,412 | 5,350 | 2,926 | 33,640 |
| Sim3 | Nanopore | 0.5M | 50 | 193 | 200 | 890 | 166 | 175 | 45 |
| Sim4 | Nanopore | 1M | 100 | 410 | 500 | 2,491 | 395 | 413 | 152 |
| Sim2021-Miseq | MiSeq | 100M | 100 | 433 | 500 | 34,211 | 3,512 | 2,014 | 25,861 |
| Sim2021-Nano | Nanopore | 1M | 100 | 433 | 500 | 2,413 | 345 | 342 | 143 |
| Gut-HiSeq | HiSeq | 53.8M | 19,053 | 20,838 | 13,903 | 130,252 | 943 | 383 | 110,128 |
| Gut-Pacbio | Pacbio | 14.7M | 19,053 | 20,838 | 13,903 | 4,671 | 64 | 8 | 723 |

Figure 2 shows the performance of PPR-Meta, viralVerify and the first phase of 3CAC on these simulated metagenome assemblies. Both PPR-Meta and viralVerify had high precision in chromosome classification, but their precision in phage and plasmid classification was usually low. Further analysis revealed that both of the algorithms distinguished well between phages and plasmids. Their low precision in phage and plasmid classification was due to contamination from chromosome contigs (Supplementary Table S1). Utilizing two-class classifiers, PlasClass and DeepVirFinder, the first phase of 3CAC improved markedly the precision in phage and plasmid classification, while it decreased a little bit the precision in chromosome classification (Figure 2, Supplementary Table S2). In contrast, recall decreased in phage and plasmid classification, but increased in chromosome classification (Supplementary Figure S1).

**Fig. 2.**
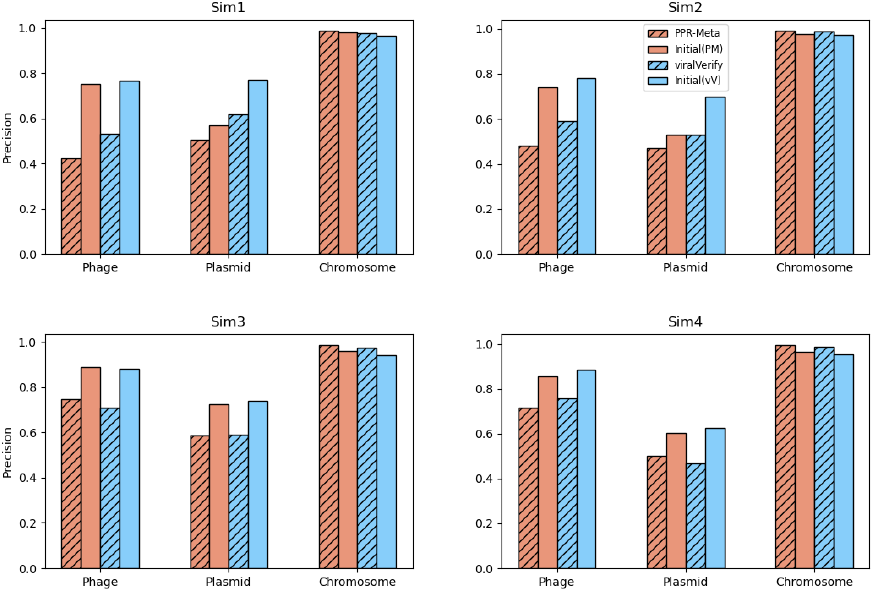
Precision of the initial classification of 3CAC compared to PPR-Meta and viralVerify. Sim1 and Sim2 are assembled from short reads. Sim3 and Sim4 are assembled from long reads. See supplementary Figure S1 for recall.

Figure 3 shows the results of initial phase of 3CAC on the short-read simulated metagenome assemblies for different contig lengths. Short contigs tended to have lower recall in the initial classification of 3CAC, while precision was not sensitive to the contig length. When the initial classification of 3CAC was generated based on PPR-Meta solution, recall decreased sharply for contigs with length *<* 1kb. When viralVerify solution was used, recall was even lower for contig shorter than 1kb and improvement with size was roughly linear. We reasoned that these classifiers classified each of the input contigs independently, and so short contigs could not be classified reliably. However, Table 1 shows that more than half of the contigs assembled from short reads are shorter than 1kb. To assist in the classification of these short contigs, 3CAC was designed to take advantage of the longer contigs with confident classification and that are neighbors of these short contigs in the assembly graph. Figure 3 shows that 3CAC significantly increased recall for all contigs with almost no loss of precision. Remarkably, the recall for contigs shorter than 1kb increased from < 0.2 to ≥ 0.8. For contigs assembled from long reads, 3CAC not only improved the recall substantially but also slightly improved the precision (Figure 4).

**Fig. 3.**
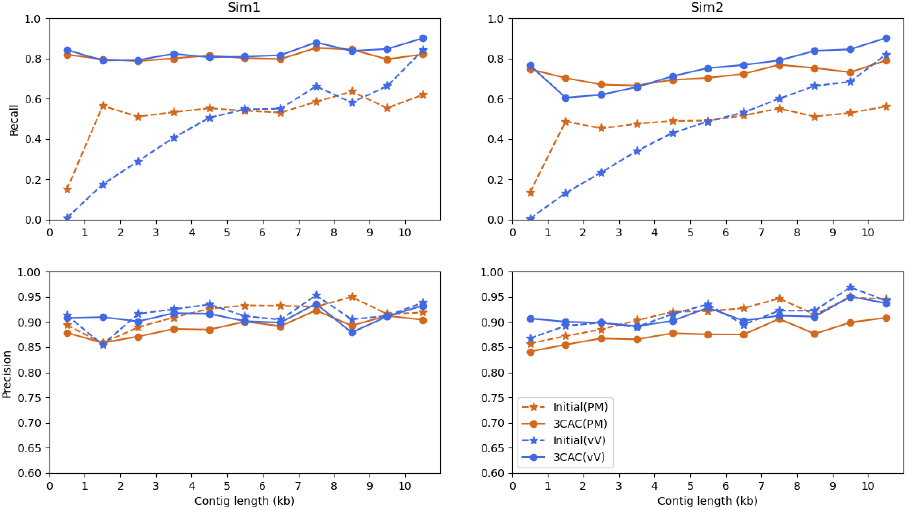
Performance on contigs assembled from simulated short reads. Results are shown for contigs of lengths < 1 kb, 1-2 kb, …,9-10 kb, ≥ 10kb.

**Fig. 4.**
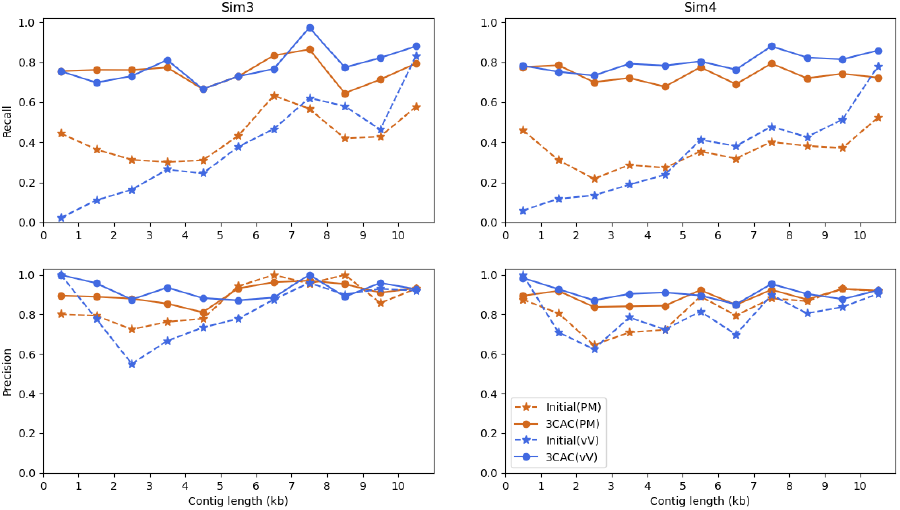
Performance on contigs assembled from simulated long reads. Results are shown for contigs of lengths < 1 kb, 1-2 kb, …,9-10 kb, ≥ 10kb.

The analysis above shows that the two phases of 3CAC algorithm improved the precision and recall for the three-class classification. Evaluation of PPR-Meta, viralVerify and 3CAC on these simulated metagenome assemblies showed that 3CAC performed the best in both precision and recall (Figure 5, Supplementary Table S1, S3). We also calculated the precision, recall and F1 scores for phage, plasmid, and chromosome classification separately (Supplementary Table S4). 3CAC had the best F1 scores on all the datasets. Note that PPR-Meta here was run with default setting. Running PPR-Meta with 0.7 score threshold (as done in Initial(PM)) resulted in higher precision but lower recall and lower F1 score. Supplementary Table S5 shows that 3CAC also outperformed PPR-Meta with 0.7 score threshold.

**Fig. 5.**
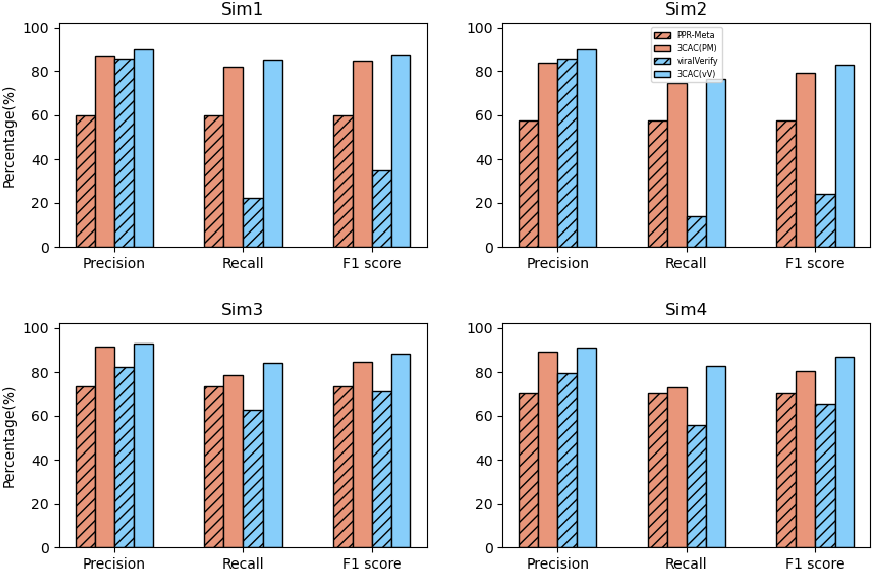
Performance of three-class classifiers on the simulated metagenome assemblies. Sim1 and Sim2 are assembled from short reads. Sim3 and Sim4 are assembled from long reads.

A potential source of bias in evaluating 3CAC is the fact that the reference genomes of our simulated metagenome assemblies may have been used for training the various classifiers. To eliminate the possible bias and evaluate the ability of 3CAC to classify novel species, additional short-read and long-read metagenome simulations were done using only genomes released on NCBI after January 2021 (Sim2021-Miseq and Sim2021-Nano in Table 1). Since all the classifiers that we used were developed prior to that date, all the tested genomes in these metagenome assemblies were not included in the training of the classifiers used by 3CAC. Evaluation on Sim2021-Miseq and Sim2021-Nano showed that 3CAC performed equally well in classification of novel species (Supplementary Figure S2 and Table S4, S5).

Overall 3CAC(vV) achieved best precision, recall, and F1 score in all six simulations, and 3CAC(PM) was a close second (Supplementary Table S5). PPR-Meta and viralVerify had 25-60 lower percentage points in F1 score on the short-read simulations, and 14-30 lower percentage points on the long-read simulations.

### 3.3 Performance on human gut microbiome samples

Five publicly available human gut microbiome samples with short-read sequencing datasets (NCBI accession numbers: ERR12976697, ERR1297651, ERR1297751, ERR1297845, ERR1297770) were selected and assembled together using metaSPAdes (Nurk *et al*., 2016). Another set of five human gut microbiome samples with long-read sequencing datasets (NCBI accession numbers: SRX2529348, SRX2529347, SRX2529346, SRX2529341, SRX2529340) were selected from Suzuki *et al*., 2019 and assembled together using metaFlye (Kolmogorov *et al*., 2020). To identify the class of contigs in the real metagenome assemblies, we downloaded all complete phage, plasmid and chromosome genomes from NCBI database and mapped contigs to all the reference genomes using minimap2 (Li, 2018). A contig was considered matched to a reference sequence if it had ≥ 80% mapping identity along ≥ 80% of the contig length. Contigs that matched to reference genomes of two or more classes were excluded to avoid ambiguity. Overall, 131,578 out of 469,022 contigs in the short-read assembly and 4,743 out of 12,541 contigs in the long-read assembly had matches to a single class and were used as the gold standard to test the classifiers. Table 1 summarizes the properties of the datasets and the assemblies.

Figures 6(a) and 7(a) show the results of PPR-Meta, viralVerify and 3CAC on the short-read and long-read assemblies, respectively. On the long-read assembly, 3CAC(vV) and 3CAC(PM) had comparable performance. 3CAC was best in precision, recall and F1 score (Figure 7). Interestingly, on the short-read assembly, 3CAC(PM) and PPR-Meta had higher F1 score than 3CAC(vV) (Figure 6 (a)). Further analysis revealed that this was due to a large number of isolated contigs in the short-read assembly graph. The second phase of 3CAC was only performed on contigs that have neighbours in the assembly graph. However, 59% of the contigs assembled from short reads were isolated and had no neighbors in the assembly graph, while the fraction on the long-read assembly was only 21%. Figures 6 (b) and 7 (b) show the results on the non-isolated contigs in the assembly graph. For both long-read and short-read assemblies, 3CAC(PM) and 3CAC(vV) had comparable performance and outperformed PPR-meta and viralVerify in precision, recall, and F1 score.

**Fig. 6.**
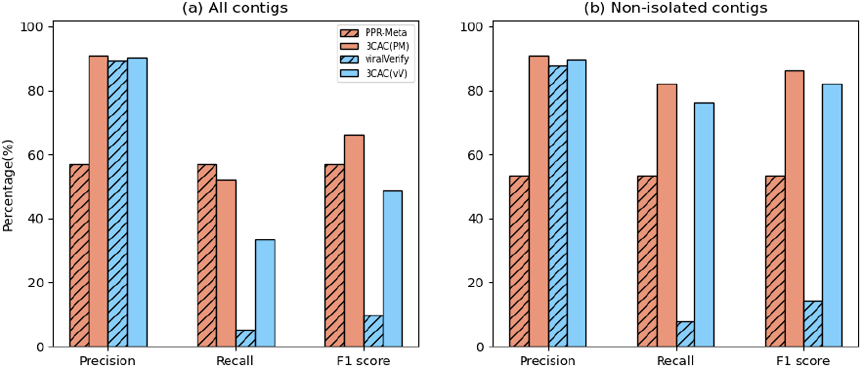
Performance of three-class classifiers on contigs assembled from short-read sequencing of human gut microbiome samples. (a) performance on all contigs; (b) performance on non-isolated contigs in the assembly graph.

**Fig. 7.**
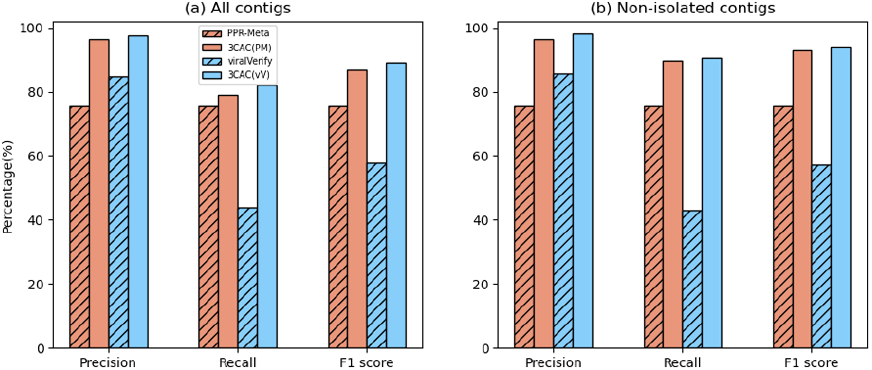
Performance of three-class classifiers on contigs assembled from long-read sequencing of human gut microbiome samples. (a) performance on all contigs; (b) performance on non-isolated contigs in the assembly graph.

Figure 8 and Supplementary Table S4 show the precision, recall and F1 score separately for phage, plasmid and chromosome classifications in both short-read and long-read assemblies. In classification of phages and plasmids, 3CAC had the highest F1 score for both short-read and long-read assemblies. The improvement in long-read assembly is more significant than in short-read assembly. This may be due to the higher quality of long-read assembly and thus the better gold standards (see more details in Discussion). PPR-Meta had the highest recall in phage and plasmid classification. However, its precision was as low as 0.01 in classifying phages from the short-read assembly, which resulted in the lowest F1 score. In chromosome classification, 3CAC had the highest F1 score in the long-read assembly while PPR-Meta performed slightly better in the short-read assembly.

**Fig. 8.**
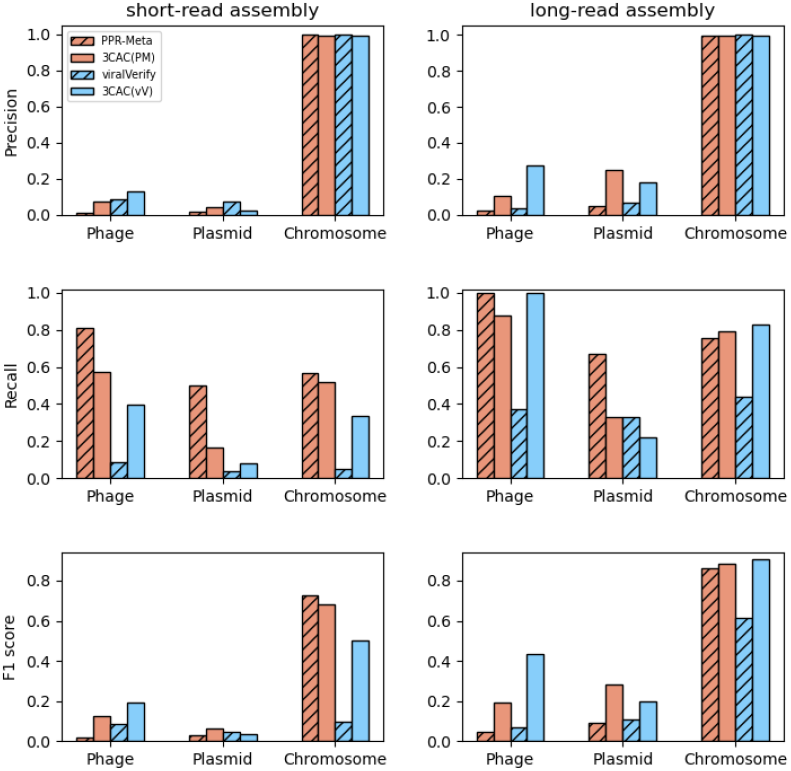
Performance of three-class classifiers in classification of phages, plasmids, and chromosomes on contigs assembled from short-read (left) and long-read (right) sequencing of human gut microbiome samples.

### 3.4 Software and Resource usage

3CAC uses classification results generated by existing classifiers as a starting point, and so the running time of its first phase depends on the classifiers used. Supplementary Table S6 shows that viralVerify was the most time consuming, followed by PPR-Meta. PlasClass and DeepVirFinder were fast as they were only run on a subset of the input contigs. Time consumed by the second phase of 3CAC was tiny compared to PPR-Meta and viralVerify. In all tests, the peak memory usage of 3CAC was less than 15 GB. Performance was measured on a 44-core, 2.2 GHz server with 792 GB of RAM. 3CAC is freely available on Github via https://github.com/Shamir-Lab/3CAC.

## 4 Discussion and Conclusion

In this study we introduced 3CAC, a new method that classifies contigs in assembly graphs into bacterial, plasmid, viral, and uncertain. 3CAC builds on initial three-class classifications generated by viralVerify or PPR-Meta and improves the classification of short and uncertain contigs by exploiting the structure of the assembly graph. Evaluation on real and simulated metagenomes assembled from both short and long reads showed that 3CAC significantly improved the recall with almost no loss of the precision. Moreover, it increased F1 score by 10-60 percentage points.

In the correction step of 3CAC, the order of treating incongruous contigs with the same number of classified neighbors may affect the results. A similar situation occurs in the propagation step. We tried several random orders and the results were very stable (< 1% difference, Supplementary Table S7).

Should a user interested specifically in a two-way classification prefer a dedicated two-way classifier over 3CAC? We compared PlasClass to 3CAC in plasmid classification, and DeepVirFinder to 3CAC in phage classification. The results on the six simulated datasets show that 3CAC had superior precision and F1 score in all cases (Supplementary Table S8 and Table S9).

Evaluation of the performance of classifiers on real metagenome assemblies remains challenging due to the lack of gold standard. By mapping contigs to all the available reference genomes, we are able to identify the class of a fraction of the contigs. However, as shown in previous studies (Fang *et al*., 2019), some plasmid genomes are quite similar to their host bacterial chromosomes. Thus, many contigs from metagenome assemblies have matches to both plasmid and chromosome reference genomes, and it is hard to identify their classes. Additionally, many contigs with no matches to the reference database may represent novel species, but they were excluded from our evaluation. Keeping in mind these shortcomings of the gold standard for real metagenome assemblies, 3CAC outperformed existing three-class classifiers substantially. The recent emergence of HiFi reads, which yield high quality of metagenome assemblies (Bickhart *et al*., 2022), may result in more complete reference databases and better gold standards for real metagenome assemblies in the future.

3CAC has some limitations. The propagation step of 3CAC can greatly improve the recall with almost no loss of the precision, but it can only be applied to non-isolated contigs in the assembly graph. The recall of isolated contigs is still limited by the performance of existing classifiers. 3CAC relies on existing two-class and three-class classifiers. In the future, we plan to turn 3CAC into a stand-alone classification tool. The detection of prophages and other non-bacterial sequences that integrate into bacterial genome is challenging to all classifiers, including 3CAC. Tools designed specifically for prophage detection are recommended for this task (Sirén *et al*., 2021; Starikova *et al*., 2020). Finally, there is room for extending 3CAC to a four-class algorithm that would be able to classify also eukaryotic contigs in metagenome assemblies (West *et al*., 2018).

## References

Antipov, D., Raiko, M., Lapidus, A., and Pevzner, P. A. (2019). Plasmid detection and assembly in genomic and metagenomic data sets. Genome Research, 29(6), 961–968.

Antipov, D., Raiko, M., Lapidus, A., and Pevzner, P. A. (2020). Metaviral SPAdes: assembly of viruses from metagenomic data. Bioinformatics, 36(14), 4126–4129.

Arredondo-Alonso, S., Willems, R. J., Van Schaik, W., and Schürch, A. C. (2017). On the (im) possibility of reconstructing plasmids from whole-genome short-read sequencing data. Microbial Genomics, 3(10).

Auslander, N., Gussow, A. B., Benler, S., Wolf, Y. I., and Koonin, E. V. (2020). Seeker: alignment-free identification of bacteriophage genomes by deep learning. Nucleic Acids Research, 48(21), e121–e121.

Barnum, T. P., Figueroa, I. A., Carlström, C. I., Lucas, L. N., Engelbrektson, A. L., and Coates, J. D. (2018). Genome-resolved metagenomics identifies genetic mobility, metabolic interactions, and unexpected diversity in perchlorate-reducing communities. The ISME Journal, 12(6), 1568–1581.

Bickhart, D. M., Kolmogorov, M., Tseng, E., Portik, D. M., Korobeynikov, A., Tolstoganov, I., Uritskiy, G., Liachko, I., Sullivan, S. T., Shin, S. B., et al. (2022). Generating lineage-resolved, complete metagenome-assembled genomes from complex microbial communities. Nature biotechnology, pages 1–9.

Calero-Cáceres, W., Ye, M., and Balcázar, J. L. (2019). Bacteriophages as environmental reservoirs of antibiotic resistance. Trends in Microbiology, 27(7), 570–577.

Fang, Z., Tan, J., Wu, S., Li, M., Xu, C., Xie, Z., and Zhu, H. (2019). PPR-Meta: a tool for identifying phages and plasmids from metagenomic fragments using deep learning. GigaScience, 8(6), giz066.

Frost, L. S., Leplae, R., Summers, A. O., and Toussaint, A. (2005). Mobile genetic elements: the agents of open source evolution. Nature Reviews Microbiology, 3(9), 722–732.

Gourlé, H., Karlsson-Lindsjö, O., Hayer, J., and Bongcam-Rudloff, E. (2019). Simulating illumina metagenomic data with insilicoseq. Bioinformatics, 35(3), 521–522.

Guo, J., Bolduc, B., Zayed, A. A., Varsani, A., Dominguez-Huerta, G., Delmont, T. O., Pratama, A. A., Gazitúa, M. C., Vik, D., Sullivan, M. B., et al. (2021). Virsorter2: a multi-classifier, expert-guided approach to detect diverse DNA and RNA viruses. Microbiome, 9(1), 1–13.

Hurwitz, B. L. and U’Ren, J. M. (2016). Viral metabolic reprogramming in marine ecosystems. Current Opinion in Microbiology, 31, 161–168.

Kieft, K., Zhou, Z., and Anantharaman, K. (2020). Vibrant: automated recovery, annotation and curation of microbial viruses, and evaluation of viral community function from genomic sequences. Microbiome, 8(1), 1–23.

Kolmogorov, M., Bickhart, D. M., Behsaz, B., Gurevich, A., Rayko, M., Shin, S. B., Kuhn, K., Yuan, J., Polevikov, E., Smith, T. P., et al. (2020). metaFlye: scalable long-read metagenome assembly using repeat graphs. Nature Methods, 17(11), 1103–1110.

Kraushaar, B., Hammerl, J., Kienol, M., Heinig, M., Sperling, N., Dinh Thanh, M., Reetz, J., Jackel, C., Fetsch, A., and Hertwig, S. (2017). Acquisition of virulence factors in livestock-associated mrsa: lysogenic conversion of cc398 strains by virulence gene-containing phages. Sci Rep 7: 2004.

Krawczyk, P. S., Lipinski, L., and Dziembowski, A. (2018). Plasflow: predicting plasmid sequences in metagenomic data using genome signatures. Nucleic Acids Research, 46(6), e35–e35.

Krishnamurthy, S. R. and Wang, D. (2017). Origins and challenges of viral dark matter. Virus Research, 239, 136–142.

Li, H. (2018). Minimap2: pairwise alignment for nucleotide sequences. Bioinformatics, 34(18), 3094–3100.

Lin, Y., Yuan, J., Kolmogorov, M., Shen, M. W., Chaisson, M., and Pevzner, P. A. (2016). Assembly of long error-prone reads using de Bruijn graphs. Proceedings of the National Academy of Sciences, 113(52), E8396–E8405.

Lopatkin, A., Meredith, H., Srimani, J., Pfeiffer, C., Durrett, R., and You, L. (2017). Persistence and reversal of plasmid-mediated antibiotic resistance. Nature Communications 8: 1689.

Mallawaarachchi, V., Wickramarachchi, A., and Lin, Y. (2020). Graphbin: refined binning of metagenomic contigs using assembly graphs. Bioinformatics, 36(11), 3307–3313.

Myers, E. W. (2005). The fragment assembly string graph. Bioinformatics, 21(suppl_2), ii79–ii85.

Nurk, S., Meleshko, D., Korobeynikov, A., and Pevzner, P. (2016). metaSPAdes: a new versatile de novo metagenomics assembler. arXiv preprint 1604.03071.

Pellow, D., Mizrahi, I., and Shamir, R. (2020). Plasclass improves plasmid sequence classification. PLoS Computational Biology, 16(4), e1007781.

Pellow, D., Zorea, A., Probst, M., Furman, O., Segal, A., Mizrahi, I., and Shamir, R. (2021). Scapp: An algorithm for improved plasmid assembly in metagenomes. Microbiome, 9(1), 1–12.

Pevzner, P. A., Tang, H., and Waterman, M. S. (2001). An Eulerian path approach to DNA fragment assembly. Proceedings of the National Academy of Sciences, 98(17), 9748–9753.

Ren, J., Ahlgren, N. A., Lu, Y. Y., Fuhrman, J. A., and Sun, F. (2017). Virfinder: a novel k-mer based tool for identifying viral sequences from assembled metagenomic data. Microbiome, 5(1), 1–20.

Ren, J., Song, K., Deng, C., Ahlgren, N. A., Fuhrman, J. A., Li, Y., Xie, X., Poplin, R., and Sun, F. (2020). Identifying viruses from metagenomic data using deep learning. Quantitative Biology, pages 1–14.

Rosenwasser, S., Ziv, C., Van Creveld, S. G., and Vardi, A. (2016). Virocell metabolism: metabolic innovations during host–virus interactions in the ocean. Trends in Microbiology, 24(10), 821–832.

Roux, S., Enault, F., Hurwitz, B. L., and Sullivan, M. B. (2015). Virsorter: mining viral signal from microbial genomic data. PeerJ, 3, e985.

Sarowska, J., Futoma-Koloch, B., Jama-Kmiecik, A., Frej-Madrzak, M., Ksiazczyk, M., Bugla-Ploskonska, G., and Choroszy-Krol, I. (2019). Virulence factors, prevalence and potential transmission of extraintestinal pathogenic escherichia coli isolated from different sources: recent reports. Gut Pathogens, 11(1), 1–16.

Simpson, J. T. and Durbin, R. (2012). Efficient de novo assembly of large genomes using compressed data structures. Genome Research, 22(3), 549–556.

Sirén, K., Millard, A., Petersen, B., Gilbert, M. T. P., Clokie, M. R., and Sicheritz-Pontén, T. (2021). Rapid discovery of novel prophages using biological feature engineering and machine learning. NAR genomics and bioinformatics, 3(1), lqaa109.

Sitaraman, R. (2018). Prokaryotic horizontal gene transfer within the human holobiont: ecological-evolutionary inferences, implications and possibilities. Microbiome, 6(1), 1–14.

Smalla, K., Jechalke, S., and Top, E. M. (2015). Plasmid detection, characterization, and ecology. Microbiology Spectrum, 3(1), 3–1.

Starikova, E. V., Tikhonova, P. O., Prianichnikov, N. A., Rands, C. M., Zdobnov, E. M., Ilina, E. N., and Govorun, V. M. (2020). Phigaro: high-throughput prophage sequence annotation. Bioinformatics, 36(12), 3882–3884.

Suzuki, Y., Nishijima, S., Furuta, Y., Yoshimura, J., Suda, W., Oshima, K., Hattori, M., and Morishita, S. (2019). Long-read metagenomic exploration of extrachromosomal mobile genetic elements in the human gut. Microbiome, 7(1), 1–16.

Thomas, C. M. and Nielsen, K. M. (2005). Mechanisms of, and barriers to, horizontal gene transfer between bacteria. Nature Reviews Microbiology, 3(9), 711–721.

Wein, T., Hülter, N., Mizrahi, I., and Dagan, T. (2019). Emergence of plasmid stability under non-selective conditions maintains antibiotic resistance. Nature Communications 10: 2595.

West, P. T., Probst, A. J., Grigoriev, I. V., Thomas, B. C., and Banfield, J. F. (2018). Genome-reconstruction for eukaryotes from complex natural microbial communities. Genome Research, 28(4), 569–580.

Yahara, K., Suzuki, M., Hirabayashi, A., Suda, W., Hattori, M., Suzuki, Y., and Okazaki, Y. (2021). Long-read metagenomics using promethion uncovers oral bacteriophages and their interaction with host bacteria. Nature Communications, 12(1), 1–12.

Yang, C., Chu, J., Warren, R. L., and Birol, I. (2017). Nanosim: nanopore sequence read simulator based on statistical characterization. GigaScience, 6(4), gix010.

Zhou, F. and Xu, Y. (2010). cBar: a computer program to distinguish plasmid-derived from chromosome-derived sequence fragments in metagenomics data. Bioinformatics, 26(16), 2051–2052.

