## supplementary tables and figures for "3CAC: improving the classification of phages and plasmids in metagenomic assemblies using assembly graphs"

#### Supplementary Material

#### A Supplementary tables

**Table A.1. Performance of viralVerify and PPR-Meta on simulated metagenome assemblies.** PPR-Meta was run with 0.7 score threshold to assure high precision.

| | True Category | # contigs | viralVerify classification | | | | PPR-Meta classification | | | |
|---|---|---|---|---|---|---|---|---|---|---|
| | | | phage | plasmid | chromosome | uncertain | phage | plasmid | chromosome | uncertain |
| Sim1 | phage | 696 | 279 | 6 | 1 | 410 | 572 | 1 | 1 | 122 |
| | plasmid | 1,699 | 23 | 380 | 58 | 1,238 | 80 | 452 | 43 | 1,124 |
| | chromosome | 12,494 | 224 | 231 | 2,624 | 9,415 | 696 | 442 | 3,749 | 7,607 |
| Sim2 | phage | 2,926 | 737 | 7 | 3 | 2,179 | 1,919 | 14 | 2 | 991 |
| | plasmid | 5,350 | 53 | 652 | 60 | 4,585 | 158 | 1,351 | 71 | 3,770 |
| | chromosome | 40,412 | 462 | 572 | 5,448 | 33,930 | 1,926 | 1,520 | 9,568 | 27,398 |
| Sim3 | phage | 175 | 163 | 2 | 0 | 10 | 164 | 0 | 0 | 11 |
| | plasmid | 166 | 9 | 124 | 12 | 21 | 1 | 88 | 5 | 72 |
| | chromosome | 890 | 58 | 84 | 485 | 263 | 55 | 62 | 366 | 407 |
| Sim4 | phage | 413 | 376 | 4 | 0 | 33 | 362 | 1 | 0 | 50 |
| | plasmid | 395 | 6 | 301 | 16 | 72 | 2 | 197 | 4 | 192 |
| | chromosome | 2,491 | 114 | 337 | 1,161 | 879 | 142 | 196 | 889 | 1,264 |
| Sim2021-Miseq | phage | 2,014 | 591 | 2 | 2 | 1,419 | 1,455 | 30 | 24 | 505 |
| | plasmid | 3,512 | 33 | 768 | 55 | 2,656 | 149 | 988 | 59 | 2,316 |
| | chromosome | 34,211 | 502 | 498 | 5,718 | 27,493 | 2,104 | 1,041 | 9,547 | 21,519 |
| Sim2021-Nano | phage | 342 | 290 | 3 | 1 | 48 | 306 | 0 | 1 | 35 |
| | plasmid | 345 | 5 | 274 | 5 | 61 | 4 | 196 | 2 | 143 |
| | chromosome | 2,413 | 153 | 215 | 1,158 | 887 | 152 | 119 | 1,045 | 1,097 |

**Table A.2. Performance of Initial(vV) and Initial(PM) on simulated metagenome assemblies.**

| | True Category | # contigs | Initial(vV) | | | | Initial(PM) | | | |
|---|---|---|---|---|---|---|---|---|---|---|
| | | | phage | plasmid | chromosome | uncertain | phage | plasmid | chromosome | uncertain |
| Sim1 | phage | 696 | 249 | 3 | 8 | 436 | 442 | 0 | 8 | 246 |
| | plasmid | 1,699 | 7 | 310 | 88 | 1,294 | 10 | 372 | 75 | 1,242 |
| | chromosome | 12,494 | 69 | 90 | 2,797 | 9,538 | 137 | 284 | 3,936 | 8,137 |
| Sim2 | phage | 2,926 | 653 | 4 | 35 | 2,234 | 1,136 | 9 | 45 | 1,736 |
| | plasmid | 5,350 | 18 | 499 | 132 | 4,701 | 33 | 964 | 213 | 4,140 |
| | chromosome | 40,412 | 165 | 213 | 5,803 | 34,231 | 363 | 842 | 10,092 | 29,115 |
| Sim3 | phage | 175 | 139 | 1 | 11 | 24 | 145 | 0 | 9 | 21 |
| | plasmid | 166 | 3 | 108 | 22 | 33 | 0 | 82 | 9 | 75 |
| | chromosome | 890 | 16 | 37 | 540 | 297 | 18 | 31 | 407 | 434 |
| Sim4 | phage | 413 | 321 | 2 | 34 | 71 | 318 | 0 | 23 | 87 |
| | plasmid | 395 | 2 | 262 | 36 | 103 | 1 | 175 | 14 | 213 |
| | chromosome | 2,491 | 38 | 153 | 1,280 | 1012 | 50 | 114 | 970 | 1,349 |
| Sim2021-Miseq | phage | 2,014 | 503 | 2 | 45 | 1,464 | 1,068 | 17 | 65 | 864 |
| | plasmid | 3,512 | 17 | 581 | 97 | 2,817 | 19 | 689 | 126 | 2,678 |
| | chromosome | 34,211 | 172 | 182 | 6,092 | 27,765 | 272 | 613 | 9,983 | 23,343 |
| Sim2021-Nano | phage | 342 | 261 | 3 | 15 | 63 | 291 | 0 | 5 | 46 |
| | plasmid | 345 | 3 | 236 | 21 | 85 | 3 | 167 | 17 | 158 |
| | chromosome | 2,413 | 36 | 96 | 1,292 | 989 | 38 | 69 | 1,129 | 1,177 |

**Table A.3. Performance of 3CAC on simulated metagenome assemblies.** 3CAC(vV) and 3CAC(PM) represent 3CAC algorithm initialized with viralVerify and PPR-Meta solutions, respectively.

| | True Category | # contigs | 3CAC(vV) | | | | 3CAC(PM) | | | |
|---|---|---|---|---|---|---|---|---|---|---|
| | | | phage | plasmid | chromosome | uncertain | phage | plasmid | chromosome | uncertain |
| Sim1 | phage | 696 | 642 | 7 | 21 | 26 | 646 | 7 | 17 | 26 |
| | plasmid | 1,699 | 18 | 1,135 | 404 | 142 | 32 | 1,103 | 411 | 153 |
| | chromosome | 12,494 | 393 | 538 | 10,901 | 662 | 434 | 902 | 10,481 | 677 |
| Sim2 | phage | 2,926 | 2,728 | 9 | 77 | 112 | 2,714 | 26 | 80 | 106 |
| | plasmid | 5,350 | 50 | 2,815 | 1,440 | 1,045 | 98 | 2,871 | 1,695 | 686 |
| | chromosome | 40,412 | 915 | 1,542 | 31,712 | 6,243 | 1,278 | 3,760 | 30,817 | 4,557 |
| Sim3 | phage | 175 | 145 | 1 | 11 | 18 | 148 | 1 | 8 | 18 |
| | plasmid | 166 | 3 | 124 | 25 | 14 | 0 | 110 | 18 | 38 |
| | chromosome | 890 | 12 | 28 | 769 | 81 | 25 | 36 | 711 | 118 |
| Sim4 | phage | 413 | 336 | 2 | 35 | 40 | 328 | 0 | 27 | 58 |
| | plasmid | 395 | 2 | 306 | 41 | 46 | 4 | 210 | 43 | 138 |
| | chromosome | 2,491 | 35 | 148 | 2,087 | 221 | 71 | 143 | 1,877 | 400 |
| Sim2021-Miseq | phage | 2,014 | 1,575 | 3 | 128 | 308 | 1,662 | 24 | 153 | 175 |
| | plasmid | 3,512 | 55 | 2,596 | 472 | 389 | 66 | 2,402 | 730 | 314 |
| | chromosome | 34,211 | 674 | 1,260 | 30,076 | 2,201 | 1,008 | 2,131 | 29,364 | 1,708 |
| Sim2021-Nano | phage | 342 | 297 | 5 | 15 | 25 | 297 | 0 | 5 | 40 |
| | plasmid | 345 | 3 | 272 | 21 | 49 | 11 | 218 | 25 | 91 |
| | chromosome | 2,413 | 33 | 94 | 2,066 | 220 | 45 | 94 | 1,961 | 313 |

**Table A.4. Performance of three-class classifiers in classification of phages, plasmids, and chromosomes on simulated and real metagenome assemblies.** PPR-Meta and PPR-Meta(0.7) represent running PPR-Meta on default setting and with a score threshold of 0.7, respectively.

| | **Tool** | **Phage** | | | **Plasmid** | | | **Chromosome** | | |
|---|---|---|---|---|---|---|---|---|---|---|
| | | **precision(%)** | **recall(%)** | **F1 score(%)** | **precision(%)** | **recall(%)** | **F1 score(%)** | **precision(%)** | **recall(%)** | **F1 score(%)** |
| Sim1 | viralVerify | 53.04 | 40.09 | 45.66 | 61.59 | 22.37 | 32.82 | 97.8 | 21.0 | 34.58 |
| | 3CAC(vV) | 60.97 | 92.24 | 73.41 | 67.56 | 66.80 | 67.18 | 96.25 | 87.25 | 91.53 |
| | PPR-Meta(0.7) | 42.43 | 82.18 | 55.97 | 50.5 | 26.6 | 34.85 | 98.84 | 30.01 | 46.04 |
| | 3CAC(PM) | 58.09 | 92.82 | 71.46 | 54.82 | 64.92 | 59.44 | 96.08 | 83.89 | 89.57 |
| | PPR-Meta | 19.06 | 94.54 | 31.72 | 25.97 | 57.74 | 35.83 | 95.31 | 58.43 | 72.45 |
| Sim2 | viralVerify | 58.87 | 25.19 | 35.28 | 52.97 | 12.19 | 19.81 | 98.86 | 13.48 | 23.73 |
| | 3CAC(vV) | 73.87 | 93.23 | 82.43 | 64.48 | 52.62 | 57.95 | 95.43 | 78.47 | 86.13 |
| | PPR-Meta (0.7) | 47.94 | 65.58 | 55.39 | 46.83 | 25.25 | 32.81 | 99.24 | 23.68 | 38.23 |
| | 3CAC(PM) | 66.36 | 92.75 | 77.37 | 43.13 | 53.66 | 47.82 | 94.55 | 76.26 | 84.43 |
| | PPR-Meta | 21.83 | 90.46 | 35.17 | 24.31 | 61.7 | 34.88 | 95.86 | 54.53 | 69.52 |
| Sim3 | viralVerify | 70.87 | 93.14 | 80.49 | 59.05 | 74.7 | 65.96 | 97.59 | 54.49 | 69.94 |
| | 3CAC(vV) | 90.63 | 82.86 | 86.57 | 81.05 | 74.70 | 77.74 | 95.53 | 86.40 | 90.74 |
| | PPR-Meta (0.7) | 74.55 | 93.71 | 83.04 | 58.67 | 53.01 | 55.70 | 98.65 | 41.12 | 58.05 |
| | 3CAC(PM) | 85.55 | 84.57 | 85.06 | 74.83 | 66.27 | 70.29 | 96.47 | 79.89 | 87.40 |
| | PPR-Meta | 60.63 | 99.43 | 75.33 | 42.04 | 84.34 | 56.11 | 97.38 | 66.85 | 79.28 |
| Sim4 | viralVerify | 75.81 | 91.04 | 82.73 | 46.88 | 76.2 | 58.05 | 98.64 | 46.61 | 63.30 |
| | 3CAC(vV) | 90.08 | 81.36 | 85.50 | 67.70 | 77.47 | 72.26 | 96.49 | 83.90 | 89.76 |
| | PPR-Meta (0.7) | 71.54 | 87.65 | 78.78 | 50.0 | 49.87 | 49.94 | 99.55 | 35.69 | 52.54 |
| | 3CAC(PM) | 81.39 | 79.42 | 80.39 | 59.49 | 53.16 | 56.15 | 96.40 | 75.35 | 84.59 |
| | PPR-Meta | 60.3 | 98.55 | 74.82 | 33.17 | 84.05 | 47.57 | 97.47 | 63.51 | 76.91 |
| Sim2021-Miseq | viralVerify | 52.49 | 29.34 | 37.64 | 60.57 | 21.87 | 32.13 | 99.01 | 16.71 | 28.60 |
| | 3CAC(vV) | 68.36 | 78.20 | 72.95 | 67.27 | 73.92 | 70.44 | 98.04 | 87.91 | 92.70 |
| | PPR-Meta (0.7) | 39.24 | 72.24 | 50.86 | 47.98 | 28.13 | 35.47 | 99.14 | 27.91 | 43.55 |
| | 3CAC(PM) | 60.75 | 82.52 | 69.98 | 52.71 | 68.39 | 59.54 | 97.08 | 85.83 | 91.11 |
| | PPR-Meta | 17.16 | 85.55 | 28.59 | 23.43 | 63.30 | 34.20 | 96.62 | 57.08 | 71.76 |
| Sim2021-Nano | viralVerify | 64.73 | 84.80 | 73.42 | 55.69 | 79.42 | 65.47 | 99.48 | 47.99 | 64.75 |
| | 3CAC(vV) | 89.19 | 86.84 | 88.00 | 73.32 | 78.84 | 75.98 | 98.29 | 85.62 | 91.52 |
| | PPR-Meta (0.7) | 66.23 | 89.47 | 76.12 | 62.22 | 56.81 | 59.39 | 99.71 | 43.31 | 60.39 |
| | 3CAC(PM) | 84.14 | 86.84 | 85.47 | 69.87 | 63.19 | 66.36 | 98.49 | 81.27 | 89.06 |
| | PPR-Meta | 36.42 | 98.28 | 53.15 | 29.39 | 87.85 | 44.04 | 99.46 | 72.45 | 83.84 |
| Gut-Hiseq | viralVerify | 8.65 | 8.88 | 8.76 | 7.13 | 3.77 | 4.93 | 99.78 | 5.15 | 9.79 |
| | 3CAC(vV) | 11.97 | 40.21 | 18.44 | 2.66 | 10.52 | 4.25 | 99.49 | 33.62 | 50.26 |
| | PPR-Meta (0.7) | 2.24 | 55.09 | 4.30 | 3.50 | 19.38 | 5.93 | 99.81 | 26.78 | 42.23 |
| | 3CAC(PM) | 7.02 | 58.22 | 12.53 | 3.75 | 16.39 | 6.11 | 99.56 | 52.15 | 68.45 |
| | PPR-Meta | 1.13 | 80.94 | 2.23 | 1.51 | 49.94 | 2.94 | 99.62 | 56.87 | 72.41 |
| Gut-Pacbio | viralVerify | 3.80 | 37.50 | 6.90 | 6.54 | 32.81 | 10.91 | 100 | 44.02 | 61.13 |
| | 3CAC(vV) | 27.59 | 100 | 43.24 | 17.65 | 23.44 | 20.13 | 99.74 | 82.89 | 90.54 |
| | PPR-Meta (0.7) | 8.25 | 100 | 15.24 | 19.55 | 40.63 | 26.40 | 99.63 | 39.94 | 57.02 |
| | 3CAC(PM) | 10.00 | 87.50 | 17.95 | 24.14 | 32.81 | 27.81 | 99.65 | 79.49 | 88.43 |
| | PPR-Meta | 2.47 | 100 | 4.82 | 4.95 | 67.19 | 9.22 | 99.55 | 75.68 | 85.99 |

**Table A.5. Performance of three-class classifiers on simulated and real metagenome assemblies.** PPR-Meta and PPR-Meta(0.7) represent running PPR-Meta on default setting and with a score threshold of 0.7, respectively.

| Dataset | Evaluation Criteria | **viralVerify** | **3CAC(vV)** | **PPR-Meta(0.7)** | **3CAC(PM)** | **PPR-Meta** |
|---|---|---|---|---|---|---|
| Sim1 | Precision | 85.81% | **90.18%** | 79.08% | 87.15% | 60.04% |
| | Recall | 22.05% | **85.15%** | 32.06% | 82.14% | 60.04% |
| | F1 score | 35.08% | **87.59%** | 45.62% | 84.57% | 60.04% |
| Sim2 | Precision | 85.53% | **90.23%** | 77.67% | 83.99% | 57.47% |
| | Recall | 14.04% | **76.52%** | 26.37% | 74.77% | 57.47% |
| | F1 score | 24.12% | **82.81%** | 39.37% | 79.11% | 57.47% |
| Sim3 | Precision | 82.39% | **92.84%** | 83.4% | 91.67% | 73.84% |
| | Recall | 62.71% | **84.32%** | 50.2% | 78.72% | 73.84% |
| | F1 score | 71.22% | **88.38%** | 62.68% | 84.70% | 73.84% |
| Sim4 | Precision | 79.4% | **91.34%** | 80.76% | 89.35% | 70.35% |
| | Recall | 55.71% | **82.81%** | 43.89% | 73.20% | 70.35% |
| | F1 score | 65.48% | **86.87%** | 56.87% | 80.47% | 70.35% |
| Sim2021-Miseq | Precision | 86.63% | **92.96%** | 77.87% | 89.05% | 59.07% |
| | Recall | 17.81% | **86.18%** | 30.17% | 84.12% | 59.07% |
| | F1 score | 29.55% | **89.45%** | 43.49% | 86.51% | 59.07% |
| Sim2021-Nano | Precision | 81.84% | **93.91%** | 84.77% | 93.22% | 74.91% |
| | Recall | 55.55% | **85.00%** | 49.90% | 79.87% | 74.91% |
| | F1 score | 66.18% | **89.23%** | 62.82% | 86.03% | 74.91% |
| Gut-Hiseq | Precision | 89.24% | 90.12% | 71.41% | **90.64%** | 56.90% |
| | Recall | 5.15% | 33.48% | 26.81% | 51.92% | **56.90%** |
| | F1 score | 9.74% | 48.83% | 38.98% | **66.02%** | 56.90% |
| Gut-Pacbio | Precision | 84.70% | **97.47%** | 90.35% | 96.35% | 75.61% |
| | Recall | 43.86% | **82.12%** | 40.05% | 78.87% | 75.61% |
| | F1 score | 57.79% | **89.14%** | 55.50% | 86.74% | 75.61% |

**Table A.6. Running time of the classifiers.** Numbers in the table show running time in minutes for each classifier. Row 3CAC shows the running time for the second phase of 3CAC algorithm.

| | **Sim1** | **Sim2** | **Sim3** | **Sim4** | **Sim2021 -Miseq** | **Sim2021 -Nano** | **Gut-HiSeq** | **Gut-Pacbio** |
|---|---|---|---|---|---|---|---|---|
| **viralVerify** | 141 | 201 | 165 | 218 | 314 | 324 | 238 | 292 |
| **PPR-Meta** | 22 | 32 | 23 | 33 | 69 | 65 | 67 | 50 |
| **PlasClass** | 0.43 | 0.61 | 0.72 | 0.72 | 0.6 | 0.68 | 0.7 | 0.22 |
| **DeepVirFinder** | 2.5 | 5.5 | 2 | 3.5 | 6.3 | 3.4 | 32 | 1 |
| **3CAC** | 1.5 | 3.5 | 0.5 | 0.5 | 2.1 | 0.5 | 4 | 55 |

**Table A.7. Performance of 3CAC with different contig scanning orders.** In the correction step, each set of incongruous contigs with the same number of classified neighbors was implemented by a queue data structure. The same was done for implied contigs in the propagation step. These queues determine the contig scanning order. To test the effect of the order, we modified 3CAC to use lists instead of queues, and inserted any new contig to a random location in the corresponding list. The table shows the results for the original 3CAC (O) and for two random orders R1 and R2.

| Dataset | Evaluation Criteria | **3CAC(vV)** | | | **3CAC(PM)** | | |
|---|---|---|---|---|---|---|---|
| | | **O** | **R1** | **R2** | **O** | **R1** | **R2** |
| Sim1 | Precision | 90.18% | 89.70% | 89.49% | 87.15% | 86.71% | 86.65% |
| | Recall | 85.15% | 84.96% | 84.79% | 82.14% | 82.05% | 82.11% |
| | F1 score | 87.59% | 87.27% | 87.08% | 84.57% | 84.31% | 84.32% |
| Sim2 | Precision | 90.23% | 90.19% | 89.76% | 83.99% | 83.72% | 83.71% |
| | Recall | 76.52% | 76.70% | 76.32% | 74.77% | 74.76% | 74.73% |
| | F1 score | 82.81% | 82.90% | 82.50% | 79.11% | 78.99% | 78.96% |
| Sim3 | Precision | 92.84% | 92.68% | 92.67% | 91.67% | 90.52% | 91.52% |
| | Recall | 84.32% | 84.32% | 84.24% | 78.72% | 78.31% | 78.88% |
| | F1 score | 88.38% | 88.30% | 88.26% | 84.70% | 83.97% | 84.73% |
| Sim4 | Precision | 91.34% | 91.09% | 90.98% | 89.35% | 89.65% | 89.32% |
| | Recall | 82.81% | 82.72% | 82.60% | 73.20% | 73.81% | 73.54% |
| | F1 score | 86.87% | 86.70% | 86.59% | 80.47% | 80.96% | 80.67% |
| Sim2021-Miseq | Precision | 92.96% | 92.92% | 92.55% | 89.05% | 88.94% | 88.73% |
| | Recall | 86.18% | 86.44% | 86.09% | 84.12% | 84.24% | 84.05% |
| | F1 score | 89.45% | 89.56% | 89.21% | 86.51% | 86.53% | 86.32% |
| Sim2021-Nano | Precision | 93.91% | 93.91% | 93.67% | 93.22% | 92.66% | 92.43% |
| | Recall | 85.00% | 85.13% | 84.90% | 79.87% | 79.45% | 79.55% |
| | F1 score | 89.23% | 89.31% | 89.07% | 86.03% | 85.55% | 85.51% |
| Gut-Hiseq | Precision | 90.12% | 89.86% | 89.73% | 90.64% | 90.50% | 90.35% |
| | Recall | 33.48% | 33.42% | 33.37% | 51.92% | 52.00% | 51.93% |
| | F1 score | 48.83% | 48.72% | 48.65% | 66.02% | 66.05% | 65.95% |
| Gut-Pacbio | Precision | 97.47% | 97.53% | 97.58% | 96.35% | 96.10% | 96.33% |
| | Recall | 82.12% | 82.24% | 82.35% | 78.87% | 78.83% | 79.00% |
| | F1 score | 89.14% | 89.23% | 89.32% | 86.74% | 86.61% | 86.81% |

**Table A.8. Comparison of PlasClass and 3CAC in plasmid classification on simulated metagenome assemblies.** PlasClass(0.5) and PlasClass(0.7) classified a contig as plasmid if its score $> 0.5$ and $\geq 0.7$, respectively. Here, precision was calculated as the fraction of correctly classified plasmid contigs among all contigs classified as plasmids, and the recall was calculated as the fraction of correctly classified plasmid contigs among all plasmid contigs.

| Dataset | Evaluation Criteria | **PlasClass(0.5)** | **PlasClass(0.7)** | **3CAC(vV)** | **3CAC(PM)** |
|---|---|---|---|---|---|
| Sim1 | Precision | 20.32% | 21.48% | **67.56%** | 54.82% |
| | Recall | **68.16%** | 57.27% | 66.80% | 64.92% |
| | F1 score | 31.30% | 31.25% | **67.18%** | 59.44% |
| Sim2 | Precision | 17.49% | 19.04% | **64.48%** | 43.13% |
| | Recall | **68.41%** | 57.42% | 52.62% | 53.66% |
| | F1 score | 27.86% | 28.60% | **57.95%** | 47.82% |
| Sim3 | Precision | 31.20% | 42.41% | **81.05%** | 74.83% |
| | Recall | **87.95%** | 82.53% | 74.70% | 66.27% |
| | F1 score | 46.06% | 56.03% | **77.74%** | 70.29% |
| Sim4 | Precision | 27.32% | 35.00% | **67.11%** | 59.49% |
| | Recall | **89.37%** | 77.97% | 77.47% | 53.16% |
| | F1 score | 41.85% | 48.31% | **71.92%** | 56.15% |
| Sim2021-Miseq | Precision | 15.67% | 16.56% | **67.27%** | 52.71% |
| | Recall | 71.50% | 58.66% | **73.92%** | 68.39% |
| | F1 score | 25.71% | 25.82% | **70.44%** | 59.54% |
| Sim2021-Nano | Precision | 26.98% | 36.40% | **73.32%** | 69.87% |
| | Recall | **86.96%** | 81.45% | 78.84% | 63.19% |
| | F1 score | 41.18% | 50.31% | **75.98%** | 66.36% |

**Table A.9. Comparison of DeepVirFinder and 3CAC in phage classification on simulated metagenome assemblies.** DeepVirFinder(0.05) and DeepVirFinder(0.03) classified a contig as phage if its pvalue $\leq 0.05$ and $\leq 0.03$, respectively. Here, precision was calculated as the fraction of correctly classified phage contigs among all contigs classified as phages, and the recall was calculated as the fraction of correctly classified phage contigs among all phage contigs. 76 contigs longer than 2Mb were excluded from the comparison since DeepVirFinder did not terminate successfully on these long contigs.

| Dataset | Evaluation Criteria | **DeepVirFinder(0.05)** | **DeepVirFinder(0.03)** | **3CAC(vV)** | **3CAC(PM)** |
|---|---|---|---|---|---|
| Sim1 | Precision | 29.46% | 39.10% | **60.97%** | 58.09% |
| | Recall | 83.76% | 77.59% | 92.24% | **92.82%** |
| | F1 score | 43.59% | 52.00% | **73.41%** | 71.46% |
| Sim2 | Precision | 30.99% | 39.52% | **73.87%** | 66.36% |
| | Recall | 70.95% | 63.81% | **93.23%** | 92.75% |
| | F1 score | 43.14% | 48.81% | **82.43%** | 77.37% |
| Sim3 | Precision | 49.09% | 60.98% | **90.63%** | 85.55% |
| | Recall | **92.00%** | 85.71% | 82.86% | 84.57% |
| | F1 score | 64.02% | 71.26% | **86.57%** | 85.06% |
| Sim4 | Precision | 46.22% | 58.63% | **90.08%** | 81.39% |
| | Recall | **90.31%** | 83.05% | 81.36% | 79.42% |
| | F1 score | 61.15% | 68.74% | **85.50%** | 80.39% |
| Sim2021-Miseq | Precision | 28.52% | 37.34% | **68.36%** | 60.75% |
| | Recall | 74.48% | 67.03% | 78.20% | **82.52%** |
| | F1 score | 47.97% | 41.24% | **72.95%** | 69.98% |
| Sim2021-Nano | Precision | 37.63% | 55.70% | **89.19%** | 84.14% |
| | Recall | **94.74%** | 90.06% | 86.84% | 86.84% |
| | F1 score | 53.87% | 68.83% | **88.00%** | 85.47% |

### B Supplementary figures

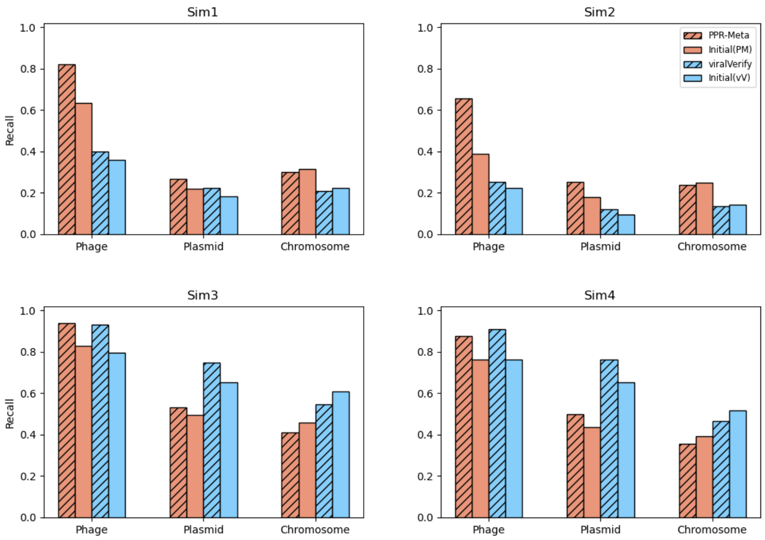


**Fig. B.1. Recall of the initial classification of 3CAC compared to PPR-Meta and viralVerify.** Sim1 and Sim2 are assembled from short reads. Sim3 and Sim4 are assembled from long reads.

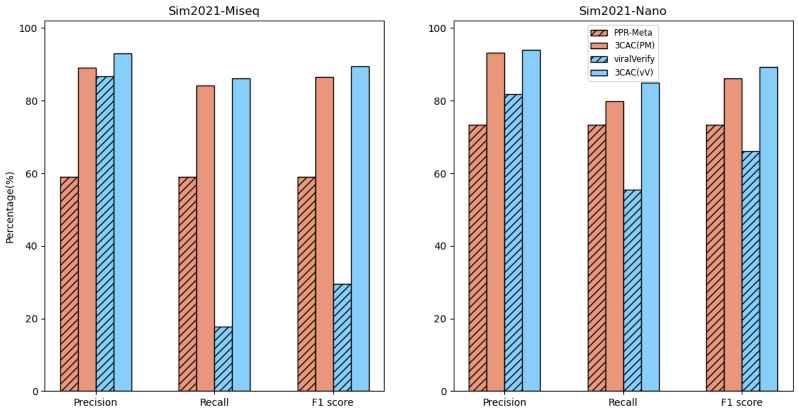


**Fig. B.2. Performance of three-class classifiers on metagenomes simulated from newly released genomes.** Performance on contigs assembled from simulated short reads (Sim2021-Miseq) and long reads (Sim2021-Nano) are shown in left and right, respectively.
